# A Mathematical Model to Simulate Intracranial Pressure and Unified Interpretation of Several Disease Entities

**DOI:** 10.1101/2020.05.31.120071

**Authors:** Jingsheng Wang, Peng Lu

## Abstract

**BACKGROUND:** Many clinical phenomena related to cerebrospinal fluid(CSF) and intracranial pressure (ICP) are often contrary to common sense and difficult to explain by classical theory. Such as slit ventricle syndrome, normal intracranial pressure hydrocephalus / low pressure hydrocephalus, paradoxical herniation, and so on. Many authors have different theories about them but can’t have an unified explanation.

**OBJECTIVE:** We try to simulate the above CSF disorders and ICP conduction with a mathematical method, and make theoretical interpretations to them.

**METHODS:** We introduced a mathematical model based on several well-accepted hypothesesto simulate human CSF physiology and propose that ICP curve should be an U-shaped curve (especially, we introduce the hypothesis that CSF also play a role of decompression). Maple software was used to draw charts according to our formula. We use the theory and intuitive charts to explain those illnesses one by one.

**RESULTS:** The formula: ICP = *μ* · *MAP* − δ · V^α^ · *μ* · *MAP* + θ · V^β^ · *μ · MAP* + *C*, and corresponding diagrams was conducted.

**CONCLUSION:** This mathematical model is a supplement to the classical Monro-Kellie’s theory, the curve and coordinate system can be used to analyze different pathophysiological states and give a reasonable unified explanation to them.

## Introduction

There is no fundamental breakthrough in the understanding of the relationship between hydrocephalus and ICP after the explanation of Monro-Kellie’s law more than 100 years ago^[1]^. In addition to ordinary hydrocephalus, slit ventricle syndrome, normal intracranial pressure hydrocephalus / low pressure hydrocephalus, widening of subarachnoid space in infants, ventricular widening / subdural effusion after bone flap decompression, paradoxical herniation after unilateral decompressive craniectomy, and so on, the complex and changeable relationship between CSF and ICP is often contrary to common sense, and clinicians have their own understanding and can not have a unified theoretical explanation^[2–3]^. This paper try to explain the above cerebrospinal fluid disorders uniformly with a mathematical model and its function diagrams.

In order to illustrate the role of cerebral blood flow pulsation, brain tissue compliance and cerebrospinal fluid pulsation in the maintenance of intracranial pressure, we designed a simplified brain model, as shown in figure 1.

**Figure 1:**
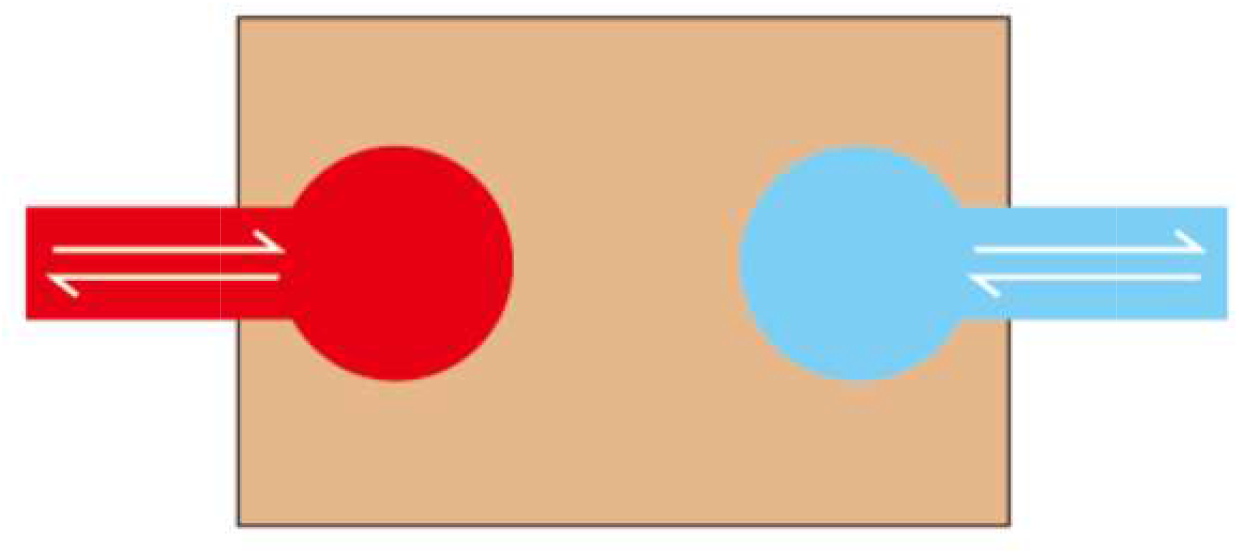
Schematic diagram of blood pulse, CSF and brain tissue: With the expansion of intracranial vascular volume (systolic blood pressure), CSF flows out of the cranial cavity; with the decrease of intracranial vascular volume (diastolic blood pressure), CSF back to the cranial cavity. Brain tissue transmits pressure between them.

### The hypothesis and the simulated mathematical model

Our theoretical derivation begins with the following hypotheses:

1. This paper assumes that the cranial cavity is a rigid container at most time, the intracranial blood vessels are elastic, and the intracranial blood volume changes cyclically with the pulsation of blood pressure;
2. The CSF in the ventricle is pushed in-and-out of the cranial cavity rhythmically under the impetus of cerebral pulsation, so as to balance the pressure in the cranial cavity.
3. The brain tissue has a certain degree of compliance, and only certain proportion of blood pulse is transmitted to the CSF in the ventricle(a linear function will be assumed in following context to simplify this relationship), causing the CSF to flow back and forth.
4. The volume of CSF in the ventricle plays a role in two aspects: on the one hand, the increase of the volume of CSF in the ventricle will increase the content of cranial cavity and hence increase ICP with marginal pressurization increasing(figure2), which is accordance with Monro-Kellie’s law; on the other hand, since CSF can flow in and out of the cranial cavity, the volume of CSF in the ventricle plays a role of buffering the transmitted blood pulse with marginal decompression decreasing(figure3). Two power functions will be assumed to simulate these relationships.
5. To simplify, we assume that ICP is mainly determined by the following 3 factors: total content within cranial cavity; the transmitted pressure from blood pulse; and decompression effect of CSF. So volume of CSF, compliance of brain tissue, patency of CSF flow, and rigidity of cranial cavity decide ICP togather. And the paper also assume a linear relationship among these factors to influence ICP.
6. We assume that the target of regulation mechanism is to select a proper volume of CSF in order to approach an proper CPP level (hence a proper ICP level, CPP=MAP-ICP, that is, the vertical distance from MAP to the curve below) by means of changing speed of absorption and producing of CSF. Besides, the regulation mechanism can be replaced or disturbed by the following factors: V-P shunt, external drainage of ventricle, and CSF absorbing dysfunction.

**Figure 2:**
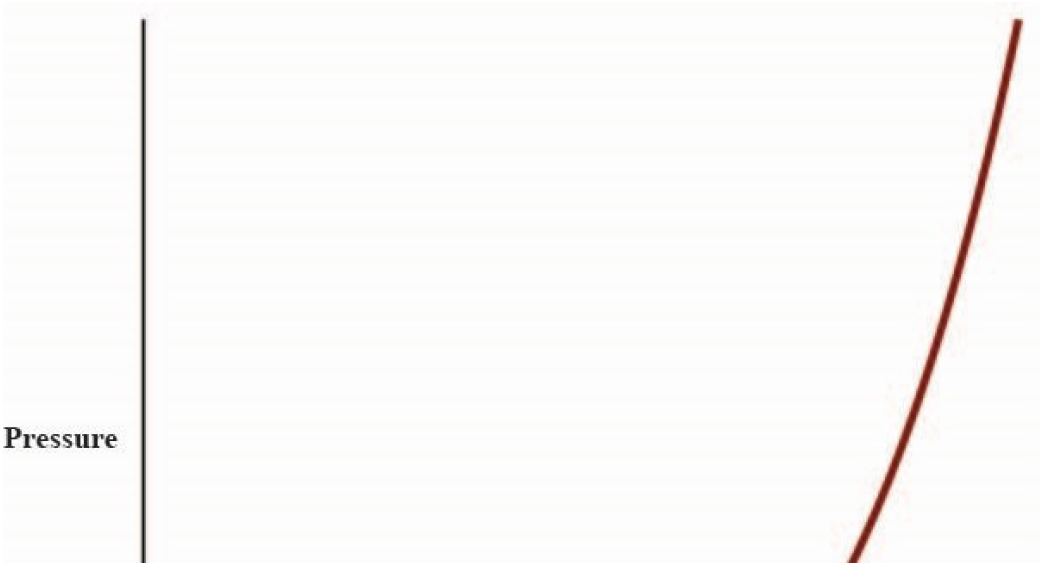
The pressurization effect of ventricle CSF with marginal pressurization increasing as V increase, which is in fact Monro-Kellie’s law.

**Figure 3:**
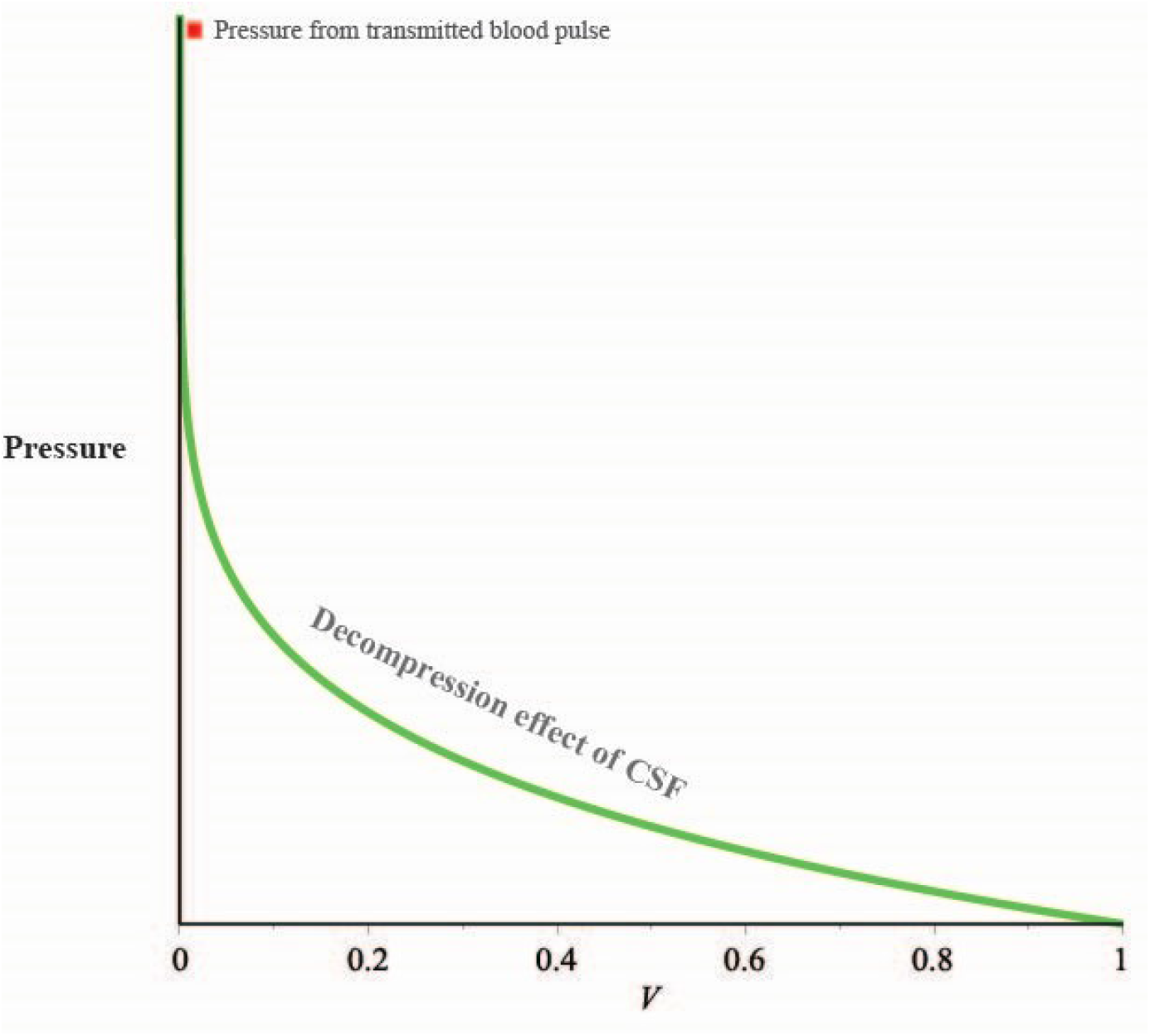
The decompression effect of ventricle CSF increases with marginal decompression decreasing as V increase.

Then we can establish a mathematical model to simulate:

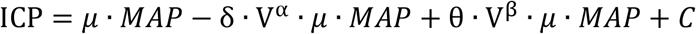

ICP: Intracranial pressure; *μ:* Compliance coefficient; MAP: Mean artery pressure; δ: Patency coefficient; V: Ventricle CSF volume; α: Depressurization constant, 0 < *α* < 1; β: Pressurization constant, β > 1; θ: Rigidity coefficient; C: Constant term。

Let the dependent variable be the ICP; independent variables include:mean artery pressure MAP; CSF volume in the ventricle V.*μ* is the brain tissue compliance coefficient with the domain of 0 < μ ≤ 1, which determines the proportion of blood pulse transmitted. *α* and β are decompression constant and pressurization constant respectively with domain of 0 < *α* < 1 (marginal decreasing as V increase) and β > 1 (marginal increasing as V increase), which influence the slop and shape of decompression and pressurization curve respectively(figure 2 and 3). δ is patency coefficient, which reflect the condition of CSF outflow tract, with domain of 0 < *δ* ≤ 1. *δ* = 1 means that the cerebrospinal fluid is unobstructed, unless obstruction or incomplete obstruction occurs, in which case 0<δ<1. θ is rigidity coefficient, which represents the rigidity of the cranial volume, with domain of θ > 0, the lower of θ the more elastic the cranial volume is, such as infants with unclosed fontanel, or patients after bone flap removal. C is a constant term which includes other factors may influence ICP in short term, such as temporarily change of cranial space by a operation or a medicine, and C will resume to the original level in long term.

### Charts of the simulated model

In order to show the above description visually, we draw the following figures respectively(curves in this paper were drawn by Maple software through substituting given values to the mathematical model to simulate different syndromes or conditions of a patient): figure2 shows pressurization of CSF which is in fact Monro-Kellie’s law; figure3 shows decompression of ventricle CSF. The combination of all of three factors can have a U-shape curve(figure 4), which simulate the relation among ICP, MAP and V.

**Figure 4:**
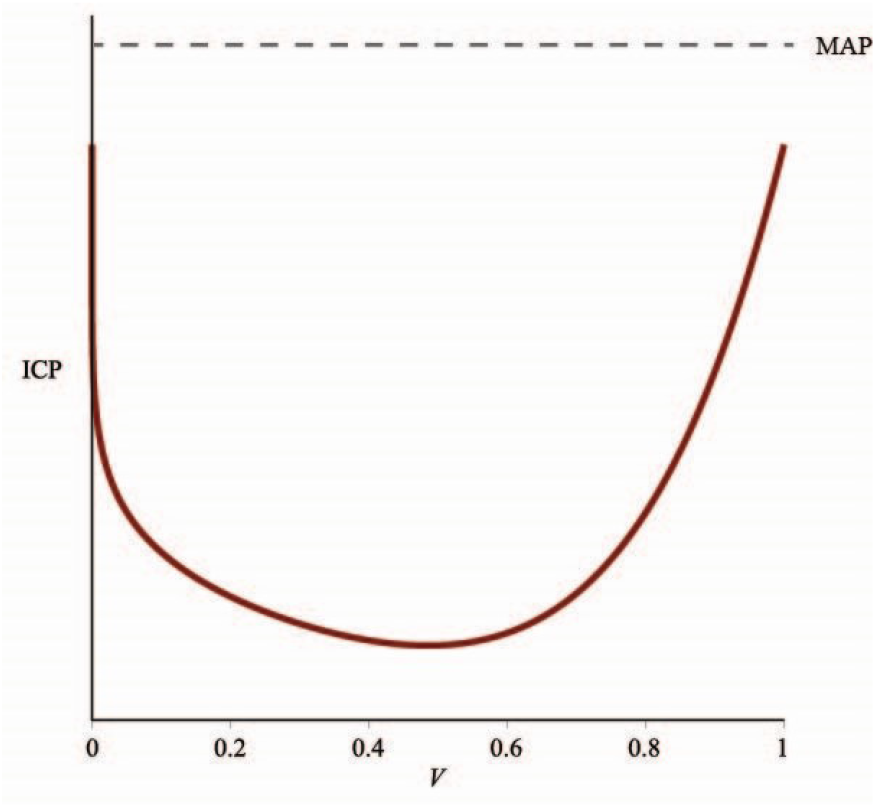
The combination of Figure2 and Figure3. this U-shaped curve directly represents our mathematically derived ICP model

Figure5 is the most important picture in this article. We can mark the position of a particular patient in this coordinate system according to the ICP and ventricle CSF volume (clinically available variables), so as to understand its pathophysiological state and help to determine treatment measures. On the upper left is the case of slit ventricle syndrome, where the ventricle CSF is low but the ICP is extremely high; on the upper right are “ordinary” hydrocephalus; on the lower right is normal pressure hydrocephalus and widening of subarachnoid space in infant which have high ventricle CSF volume and normal or even low ICP; the bottom of the middle U-shaped curve represents the “comfort zone” of normal ICP and normal ventricle size. But we should make some supplement: in the derivation of this model, the zero point of ICP is at the lowest point of the cranial cavity, which is different from the usual set at the midpoint of bilateral external auditory meatus, so there is no negative intracranial pressure theoretically, which is about 8cmH2O higher than that measured by common methods(radius of the cranial cavity).

**Figure 5:**
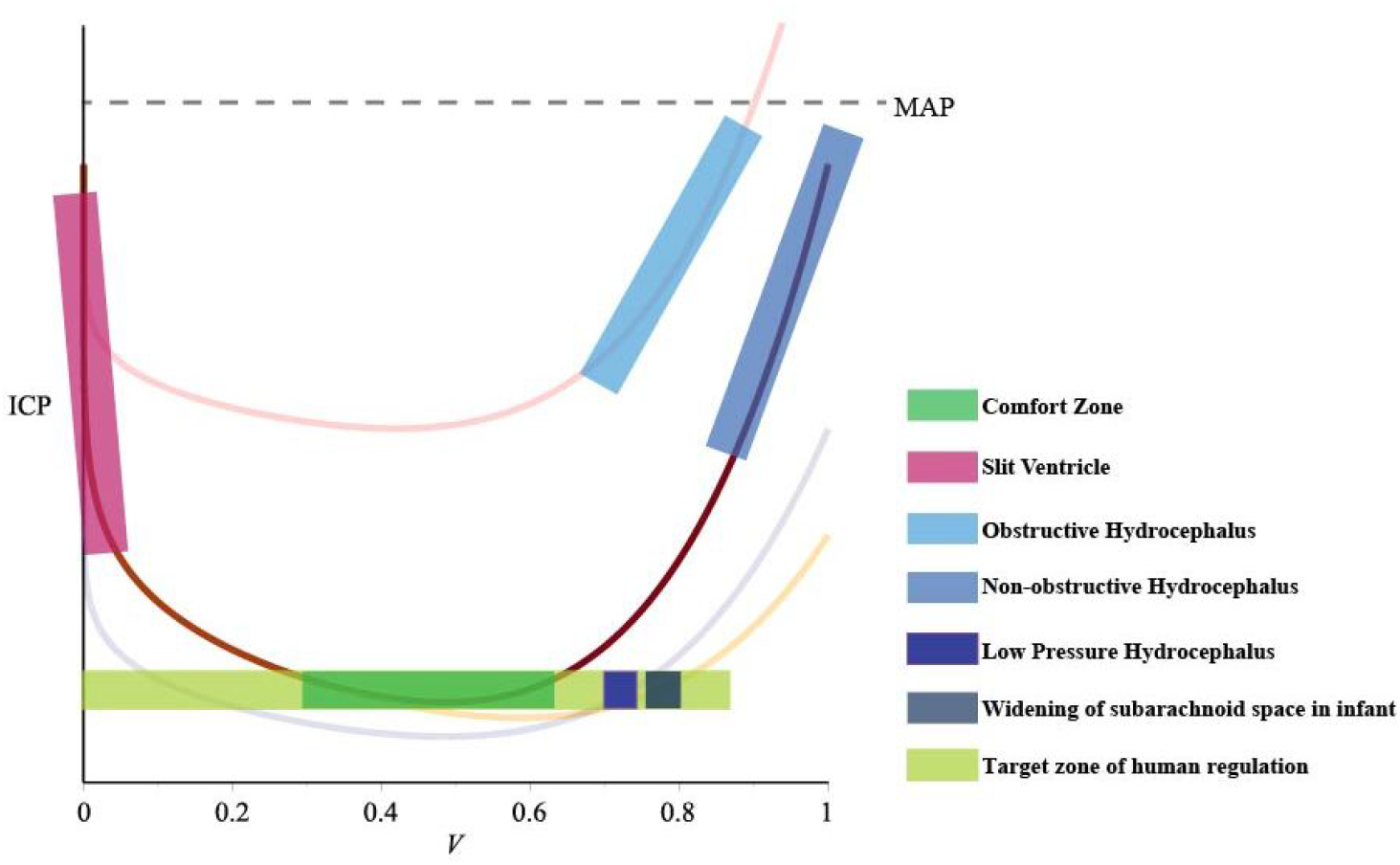
The ICP curves of different syndromes, which shift to different directions when coefficients of the model have changed. Different regions representing different syndromes.

To sum up, we can draw a “spectrum” map to mark a variety of normal and abnormal ICP-CSF states on a plane coordinate system with ICP as the longitudinal axis and ventricle CSF volume as the transverse axis. On the map there include normal health state(green comfort zone at the bottom of the U-shaped curve), hydrocephalus, slit ventricle syndrome, normal intracranial pressure hydrocephalus / low pressure hydrocephalus, infant subarachnoid space widening, ventricular widening / subdural effusion after bone flap decompression, paradoxical herniation after unilateral decompressive craniectomy, and so on. There is no clear boundary between the above adjacent marked areas, such as the subdural effusion after bone flap decompression and the widening of infant subarachnoid space, but it can be distinguished by clinical features easily.

Afterwards we will explain the applications of the model to each syndromes respectively in the following argument.

### The application and pathological interpretation of the mathematic model

#### Obstructive hydrocephalus / non-obstructive hydrocephalus

The main essence of hydrocephalus is excessive ventricle CSF and elevated ICP. When V in the formula is larger than “comfort zone”, it causes ICP to rise, and the position on the U-shaped curve moves to the right branch(Figure6). But obstructive hydrocephalus and non-obstructive hydrocephalus are different. In obstructive hydrocephalus, outflow tract was blocked, which render the patency coefficient δ decrease (δ ↓), and the curve shift towards upper left (Figure 6). In the meantime, CSF regulation mechanism is out of function since outflow tract was blocked. The volume of CSF is always increasing until ICP reach *μ*MAP. Hence, the zone of obstructive hydrocephalus locates on the upper right branch of the new curve. While in non-obstructive hydrocephalus, CSF regulation mechanism doesn’t work, and ICP increases with the volume of CSF, the patency coefficient δ remains unchanged (Figure 6).

**Figure 6.**
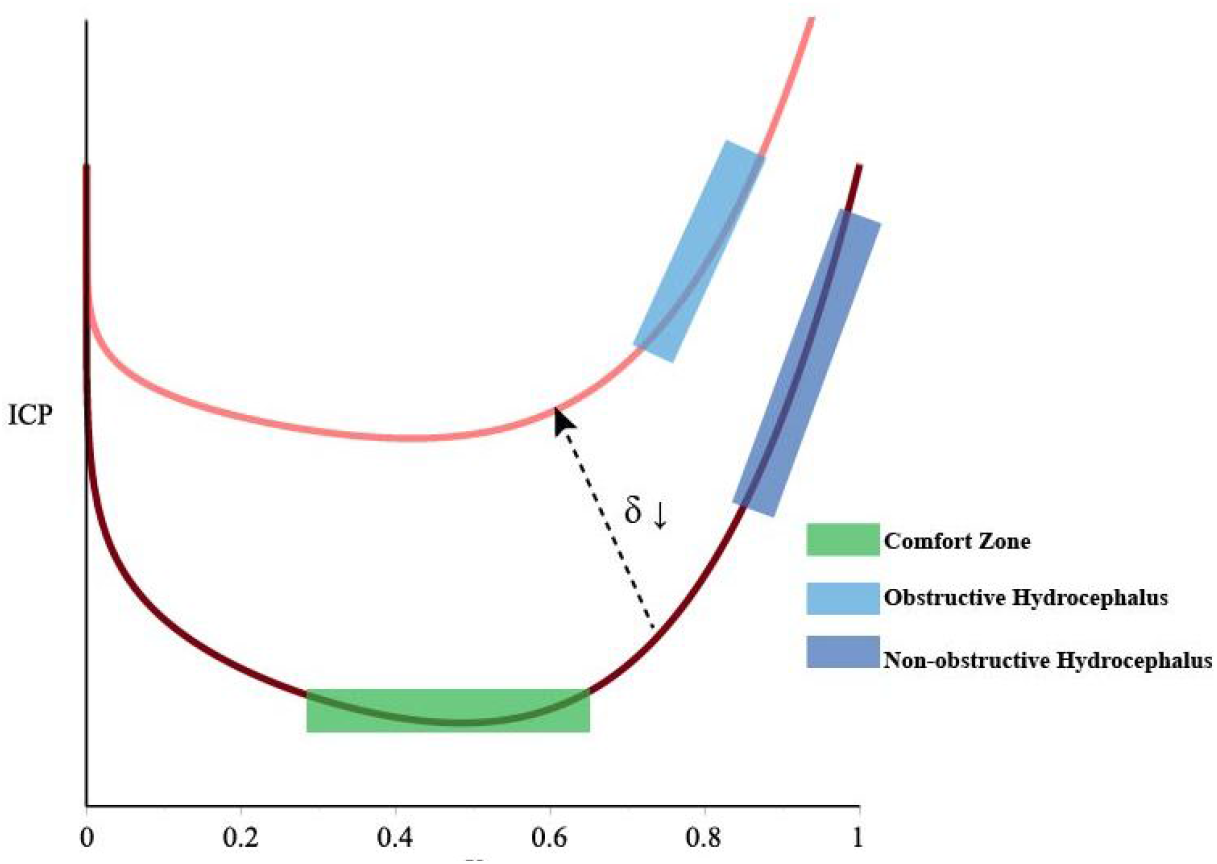
Obstructive hydrocephalus / non-oostructive hydrocephalus

After treatment of obstructive hydrocephalus such as V-P shunt or ETV^[4–5]^, patency coefficient increase (δ ↑) since CSF can be released by the shunt, and the curve shift back to original position. V will be reduced by shunt so ICP can approach to the comfort zone. After V-P shunt treatment of non-obstructive hydrocephalus, ICP decreases with the volume of CSF (figure 7).

**Figure 7.**
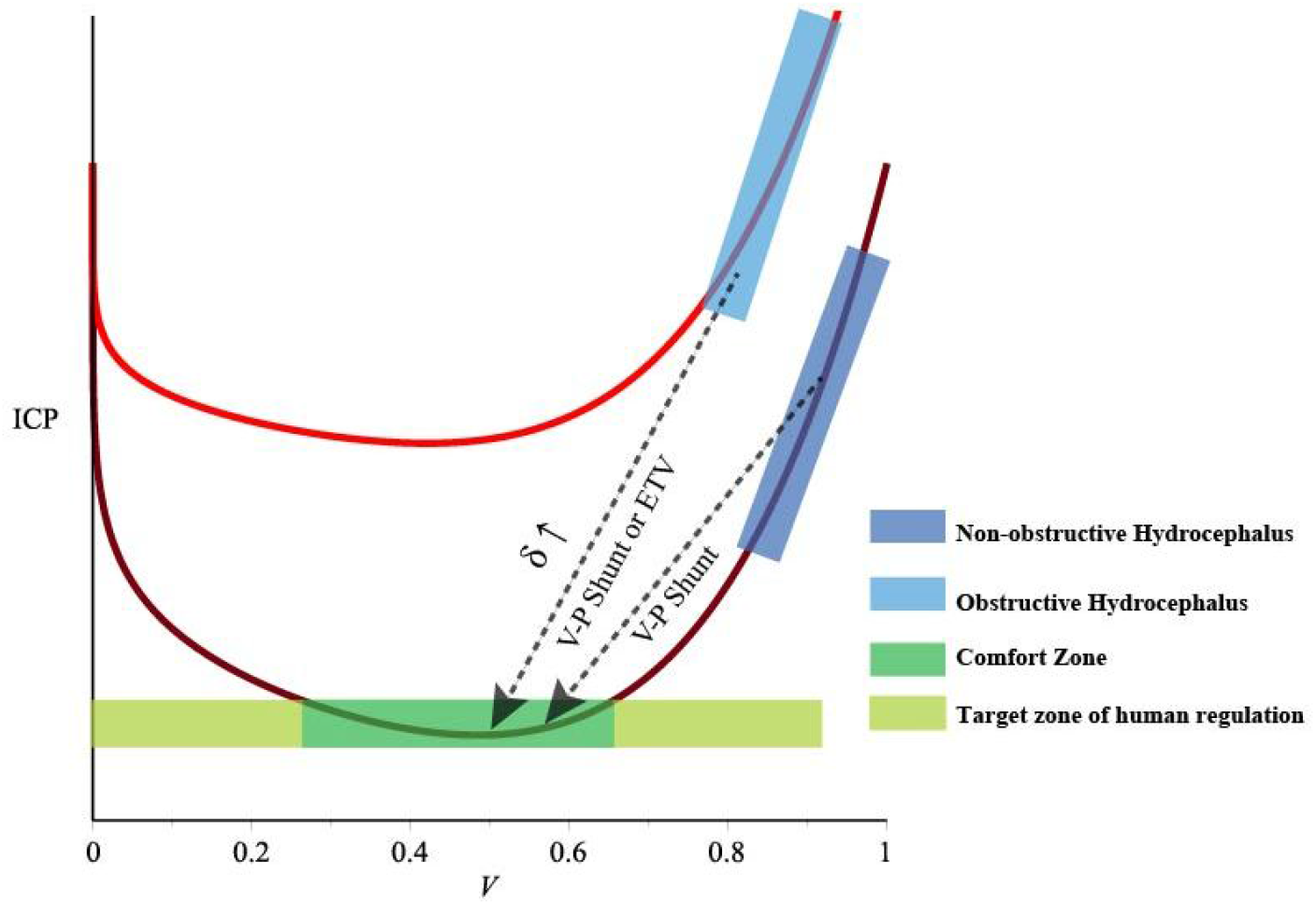

#### Slit ventricle Syndrome

Slit ventricular syndrome occurs after V-P shunt, with excessive reduction of the ventricle and elevated ICP accompanied by severe headache, optic papilla edema and other symptoms of high ICP, typically observed in older hydrocephalic children operated on in early infancy. In some literature it has been reported that ICP can rise to more than 1000mmH_2_O, which cannot be explained by classical Monro-Kellie’s law^[6–7]^. In our model it is the explanation that decrease of V leads to the rise of ICP, that is, the curve is located on the left branch of the curve (Figure8).

**Figure 8.**
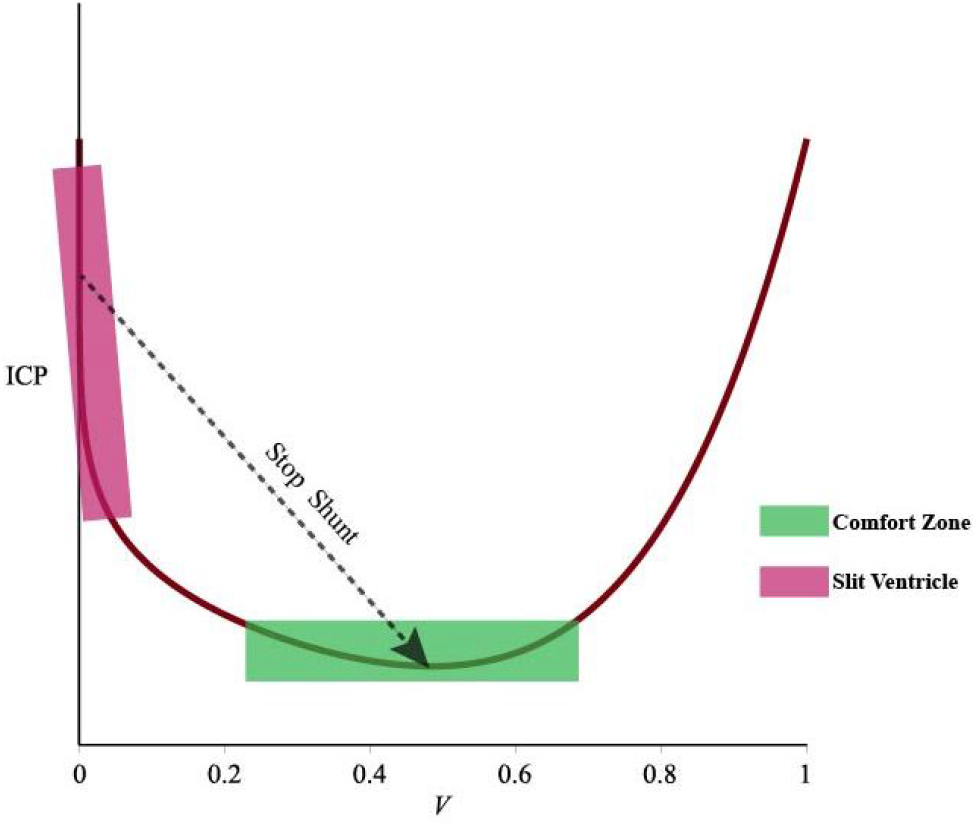

We believe that the cause of SVS is CSF regulation mechanism failure because of excessive shunt. In the acute phase, a small volume of CSF loses the ability to buffer intracranial arterial pulsation, and the ICP tends to be close to *μ*MAP. Elevated ICP then further leads to excessive shunt of CSF. Chronic SVS leads to a decrease in brain tissue compliance (*μ*↓), the curve shifts down, resulting in a decrease in ICP (Figure9).

**Figure 9.**
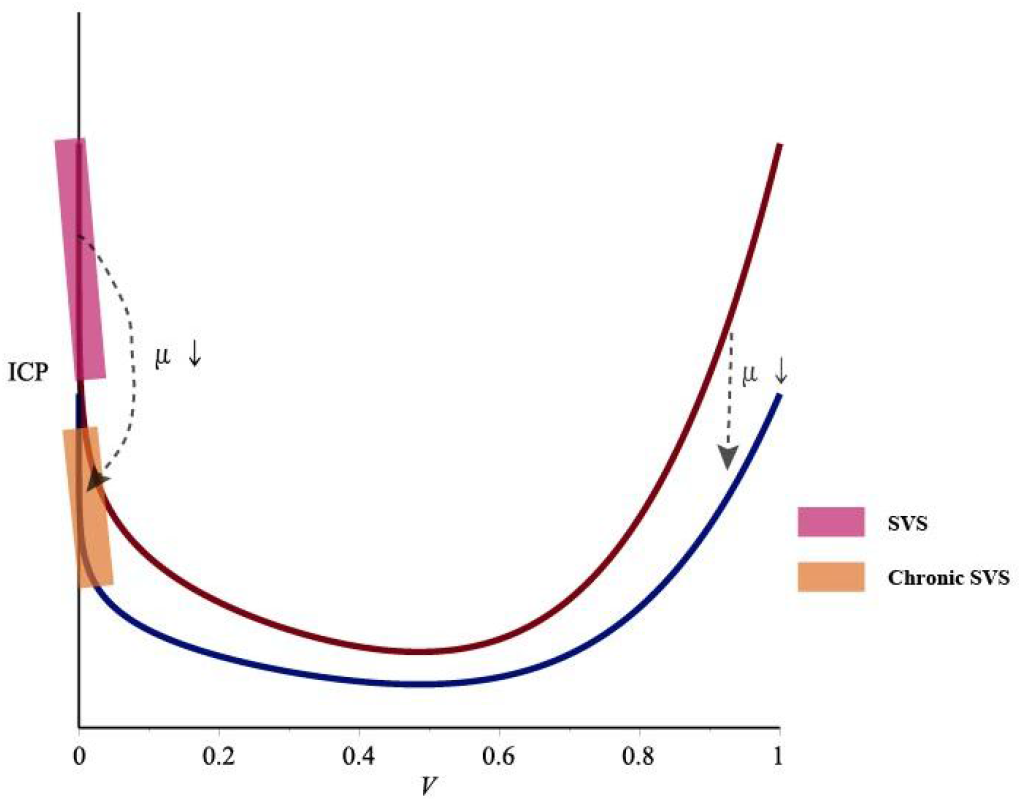
SVS

The primary treatment is to reduce excessive ICP while restoring the buffering capacity of CSF. Close the shunt valve helps to move back to the “comfort zone”, for on the left branch of curve increasing CSF causes lower ICP, contrary to common sense. The brain tissue compliance *μ* takes a relatively long time to recover afterwards.

It has also been reported that the use of subtemporal muscle decompression and cranial vault expansion can relieve slit ventricular syndrome, which can be explained that the addition of extra cranial space(C↓) and rigidity of the cranial volume (θ↓) shift the U-shaped curve to the lower right(figure10), thus reducing ICP while CSF relatively unchanged^[8–9]^.

**Figure 10.**
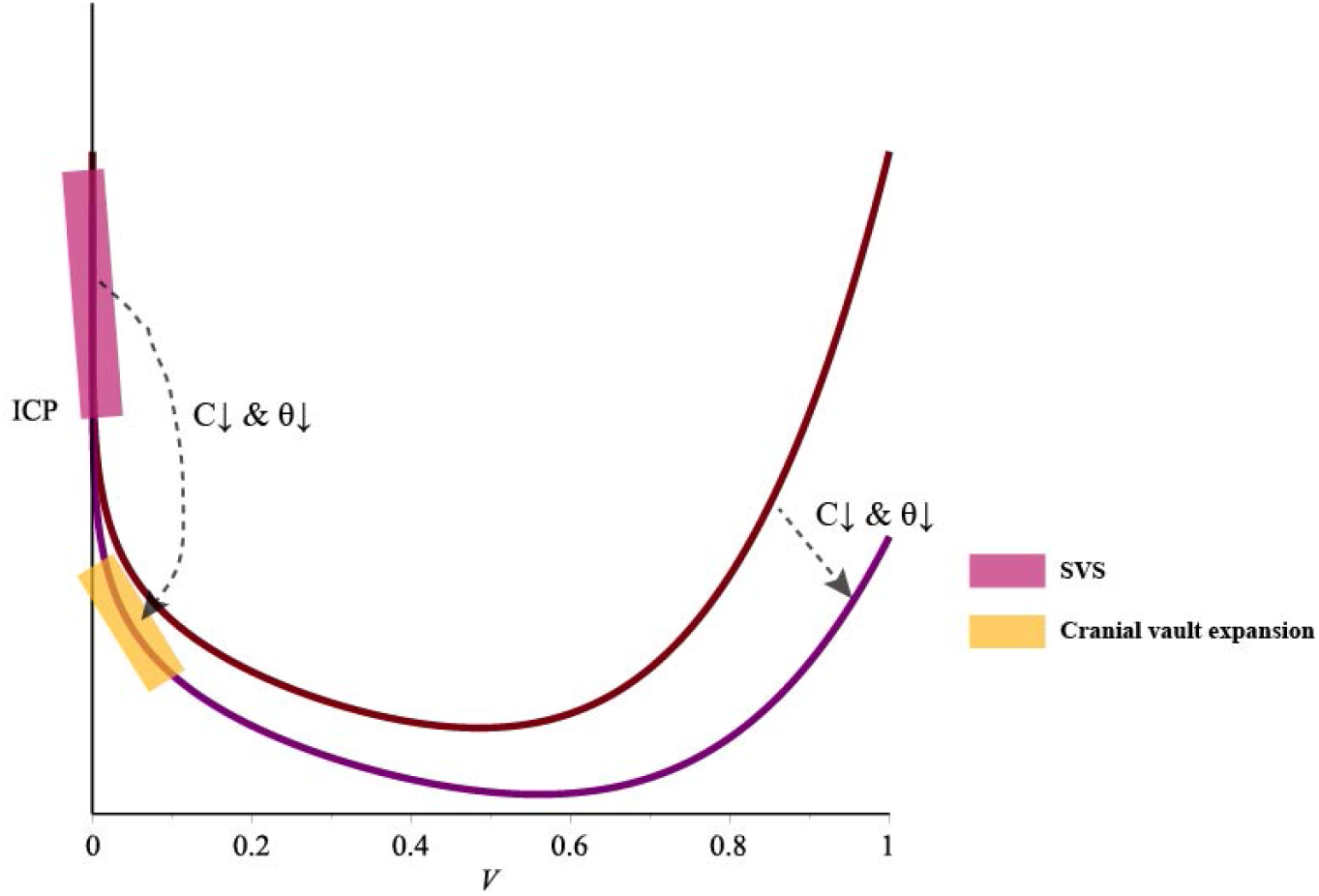

#### Normal pressure hydrocephalus / low pressure hydrocephalus

This is a very thorny subject for clinical authors and researchers. It is often seen in the elderly and occasionally in young people and adolescents. It is generally believed to be caused by decreased brain compliance, with use of ultra-low pressure drainages treatment, such as external drainage pressure lower than external auditory canal connection, ventriculo-thoracic shunt, straight tube, etc^[10]^. But there are different complex models to explain its mechanism^[11–12]^. We believe that the occurrence of this disease is due to the decrease of brain tissue compliance (*μ*↓) caused by some pathological factors such as craniocerebral trauma, subarachnoid hemorrhage, intracranial infection, encephalitis, etc. which leads to the curve moving down as a whole(Figure11). Low subarachnoid pressure leads to slower absorption of CSF (the regulation mechanism of human body) resulting in hydrocephalus (V↑). The location on the map tends to be on the right of “comfort zone” thus keeping CPP constant. The main treatment is drainage of CSF (V↓), and then wait for the natural recovery of brain tissue compliance (*μ*↑).

**Figure 11.**
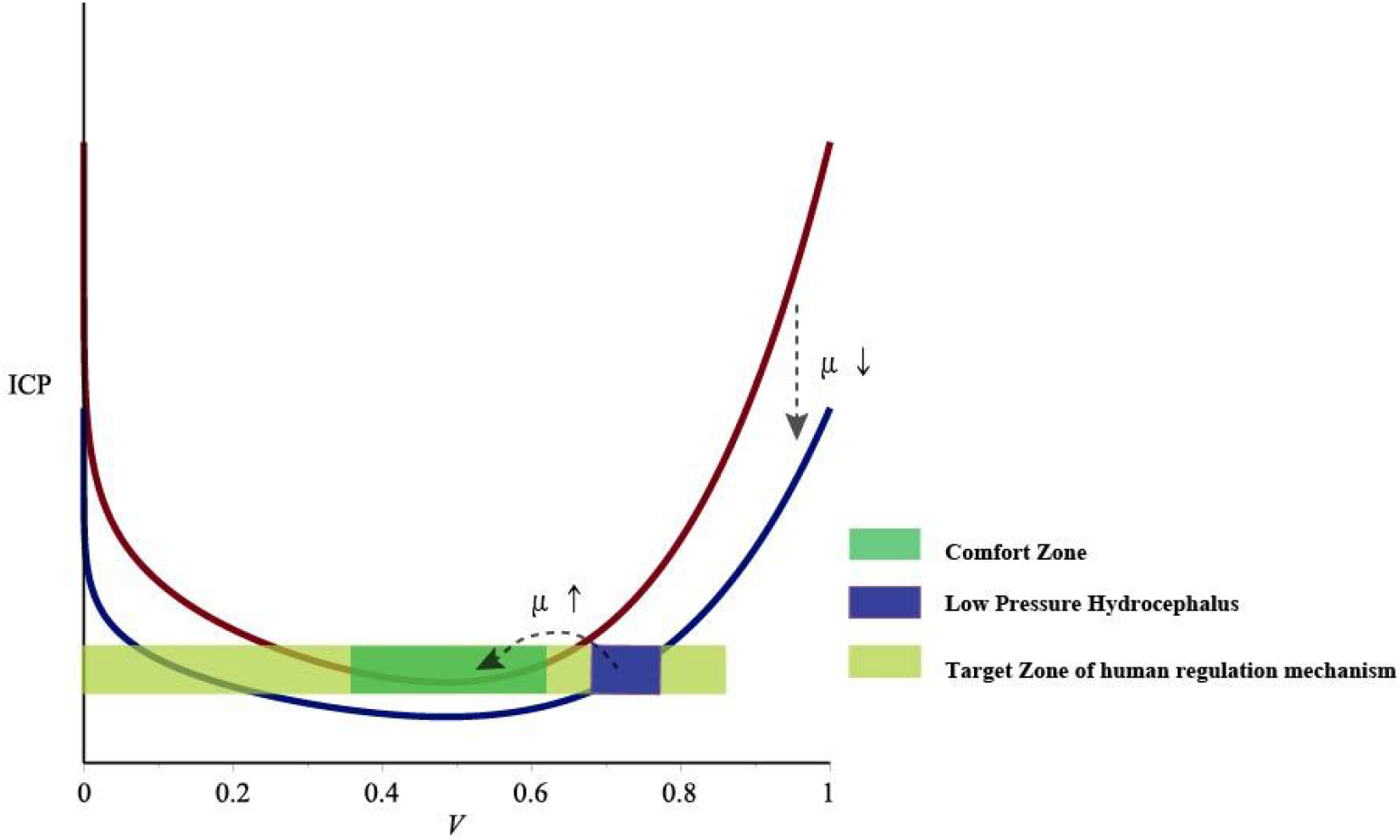
normal pressure hydrocephalus

#### Widening of subarachnoid space in infant

Widening of subarachnoid space in infants is common in infants with unclosed fontanel, children without abnormal clinical manifestations, CT/MRI subarachnoid space widening, usually without special treatment, its clinical significance is small, but the occurrence mechanism can be reasonably explained in this model. It is because when the fontanel is not closed, the non-rigid cranial cavity has a buffering effect, which increases the intracranial compensatory ability (θ↓), causing the curve shift toward the lower right (Figure12).

**Figure 12.**
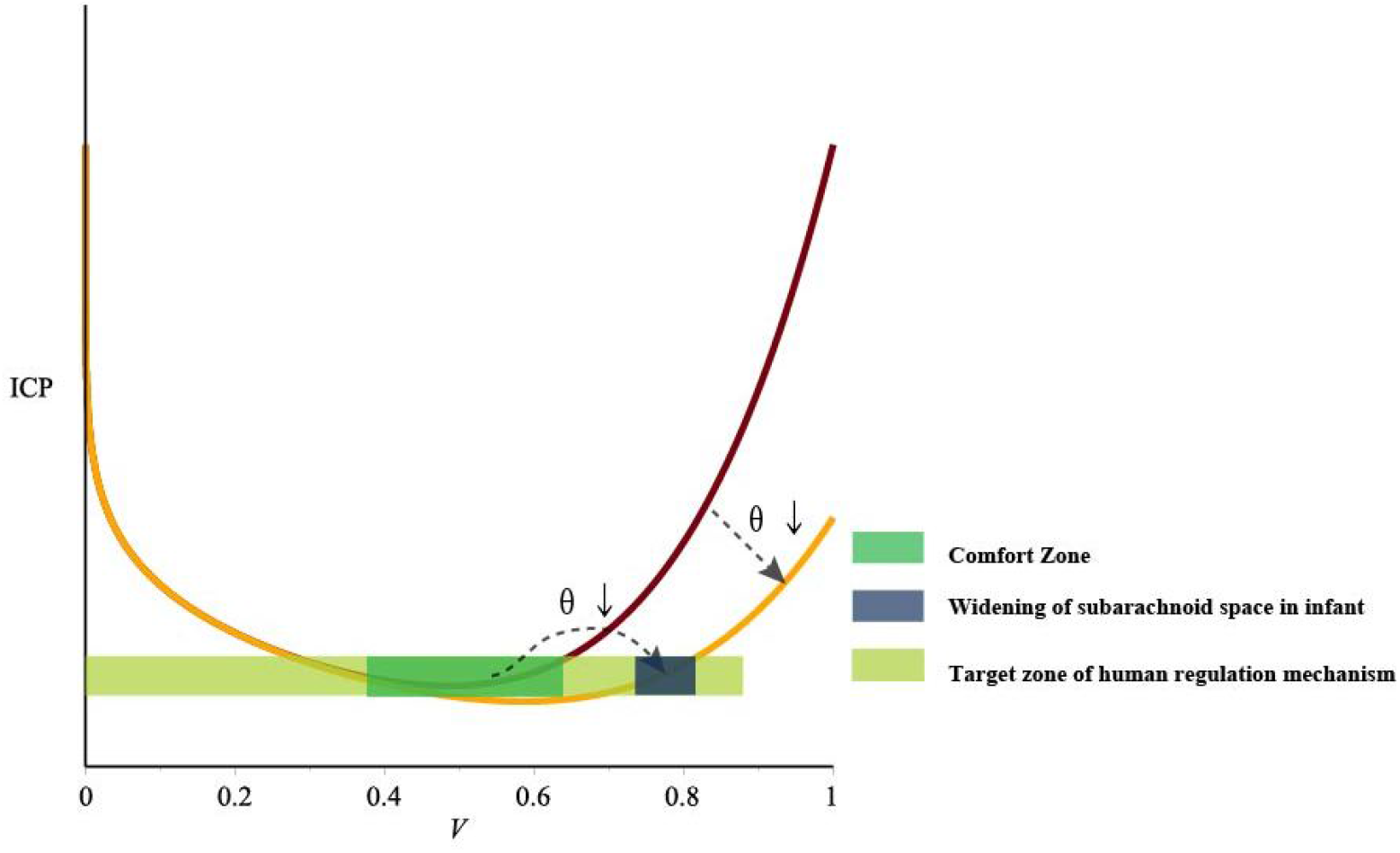
widening of subarachnoid space in infant

#### Ventricular broadening / subdural effusion after decompressive craniectomy

Ventricular broadening / subdural effusion is often observed after decompressive craniectomy, which usually relieves after bone flap repair spontaneously and requires few additional clinical treatment. However, in severe cases, temporary lumbar puncture drainage is needed to alleviate the disease^[13]^. Its mechanism can be explained by the model: due to the increase of compensatory capacity and reduction of rigidity (C↓ & θ↓) after decompressive craniectomy, the curve shift toward the lower right (figure13). According to the shifted curve, ICP slump (ICP↓) when V remain temporarily unchanged, which is out of comfort zone. Then, CSF increases(V↑) slowly by the regulation mechanism in order to approach target zone.

**Figure 13.**
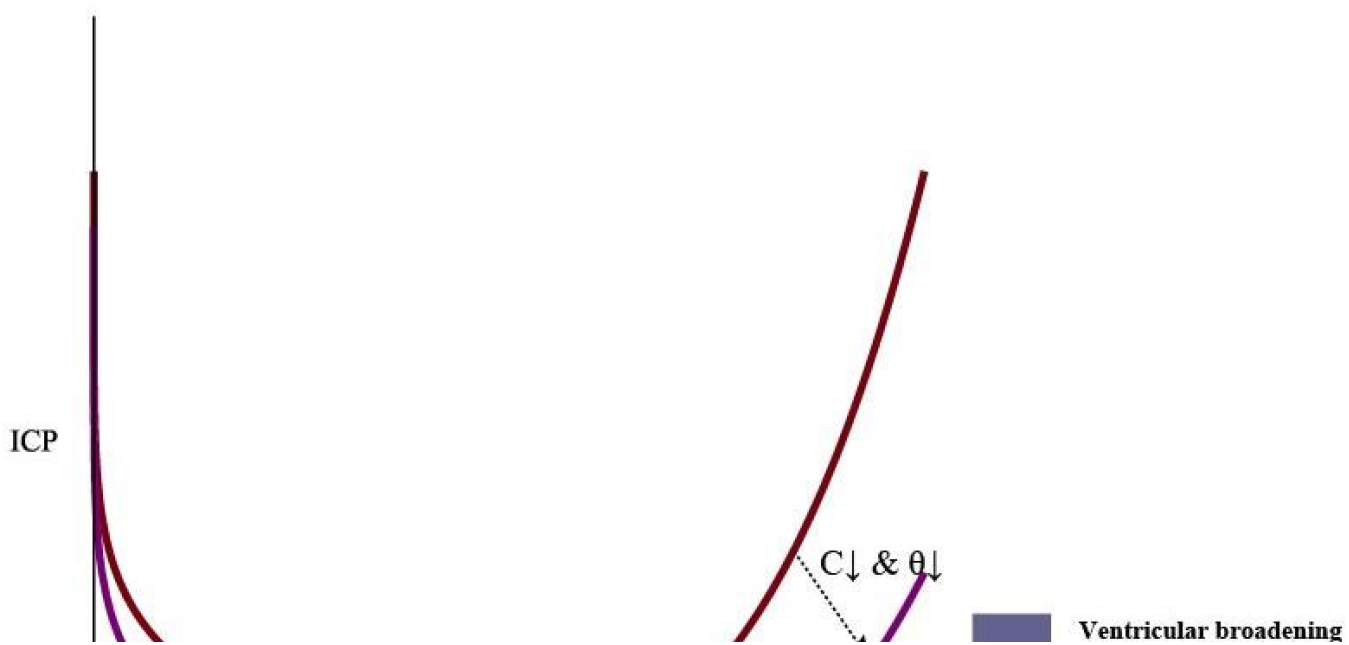

#### Paradoxical herniation

Paradoxical herniation after unilateral decompressive craniectomy is a rare and serious complication, occurs after brain swelling subsided, and inappropriate CSF drainage performed^[14]^. It has clinical importance and some effective treatments yet, but lacks a unified theoretical explanation. After decompressive craniectomy, the curve shifted toward the lower right(C↓ &θ↓), if improper CSF drainage is performed (C further decreases), ICP will go even under the atmospheric pressure (figure 14), so brain tissue is pressed inward and forms herniation. The cure can be easily understood: elevating ICP by fluid resuscitation and Trendelenburg positioning immediately, and cranioplasty afterwards.

**Figure 14.**
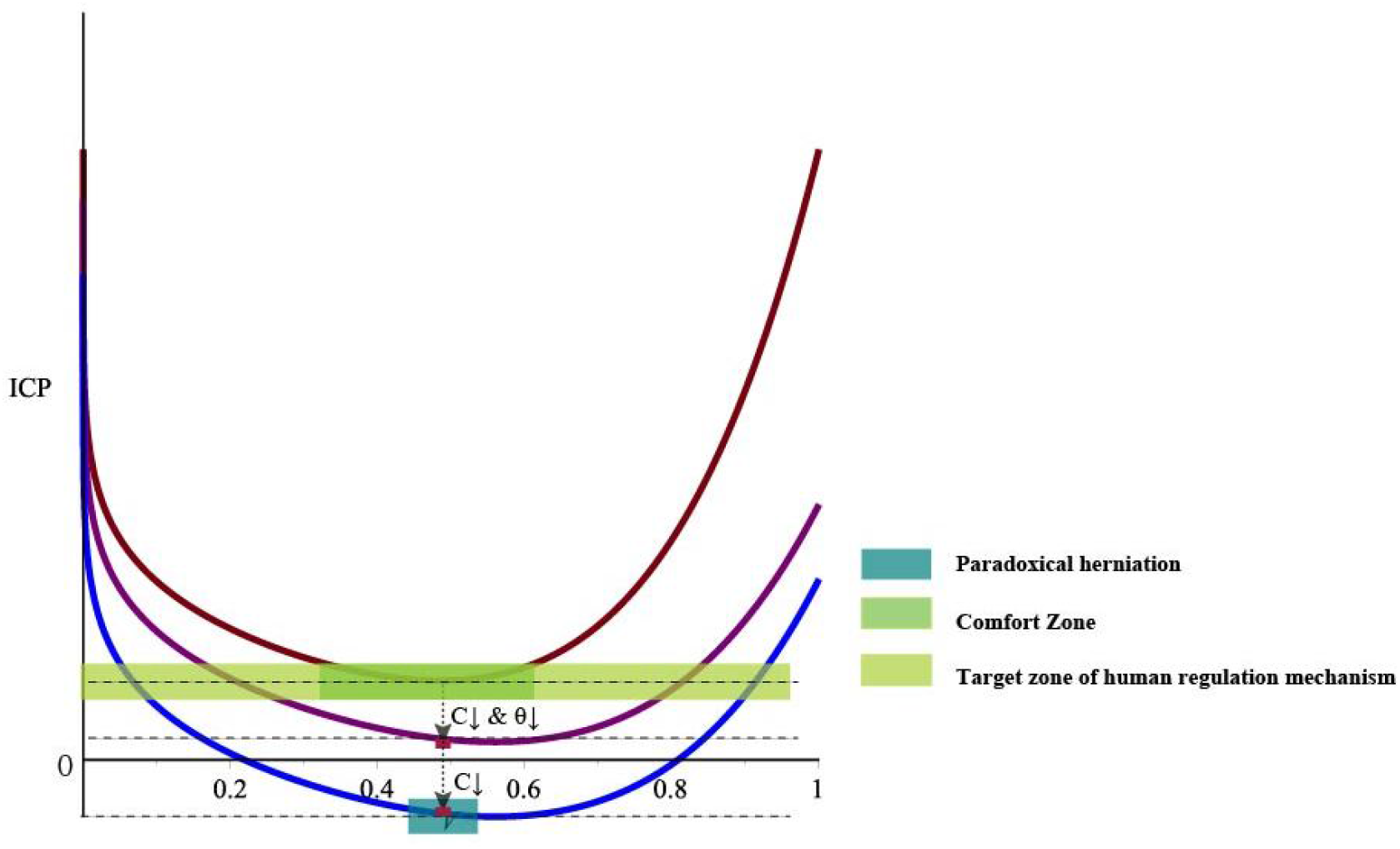

### Human regulation mechanism

In most cases the regulation of ICP is mainly on change of CSF volume: higher ICP tends to produce less and absorb more CSF; lower ICP tends toproduce more and absorb less CSF, so as to come back to “comfort zone”. Usually ICP remains in a steady state.In rare situations such as SVS, CSF volume goes to excessive low hence loses its buffering capacity, ICP pulsatile peak comes to very high level. Contrary to our common sense, if valve pressure level is then elevated, increased CSF will decrease ICP. Besides from CSF volume, compliance of brain tissue, patency of CSF flow, and rigidity of cranial cavity can also influence ICP in different manners.

## Conclusion

To sum up, for many clinical phenomena related to CSF are often contrary to common sense and difficult to explain by classical theory, we introduce a mathematical simulationand make an intuitive U-shaped curveto answer those questions based on some well-accepted hypotheses. This mathematical model is a supplement to the classical Monro-Kellie’s theory. But this study does not consider dynamic changes of blood pressure and ICP, which may influence treatment option such as V-P shunt or ETV, will be further elaborated in the next study.

